# Oncoprotein GT198 is a direct target of taxol

**DOI:** 10.1101/675579

**Authors:** Zheqiong Yang, Vadim J. Gurvich, Mohan L. Gupta, Nahid F. Mivechi, Lan Ko

**Author notes:** Address correspondence to: Lan Ko, MD/PhD, Augusta, GA, USA.

## Abstract

Taxol (paclitaxel) is one of the most successful chemotherapeutic drugs in the treatment of human cancer. It has recently been questioned whether the mechanism of action in mitotic arrest, which is ubiquitously present in all cells, is sufficient to explain the tumor specificity, clinical efficacy, and side effects of taxol. In this report, we have identified a new protein target of taxol as GT198 (gene symbol *PSMC3IP*, also known as Hop2). GT198 is an oncoprotein and a DNA repair factor involved in human common solid tumors. The *GT198* gene carries germline mutations in breast and ovarian cancer families and recurrent somatic mutations in tumor microenvironment. Mutant GT198 was identified in pericyte stem cells on capillary blood vessels inducing tumor angiogenesis. GT198 is a DNA-binding protein dimer, also stimulates DNA repair, regulates meiosis, participates in homologous DNA recombination, and activates nuclear receptor-mediated gene expression. Here we show that taxol directly binds to the DNA-binding domain of GT198 *in vitro*. Taxol serves as an allosteric inhibitor to block DNA binding to GT198 with an IC50 of 8.6 nM. Labeled taxol colocalizes with GT198 in interphase nuclei of cultured cells. Decreased GT198 expression desensitizes taxol-induced cell death, and taxol inhibits GT198 nuclear foci formation during DNA repair. Together, these results demonstrate that GT198 is a previously unrecognized direct protein target of taxol. The finding of taxol target as an oncoprotein GT198 in common solid tumors provides a rationale for the clinical efficacy of taxol. We anticipate that GT198 may serve as a clinical predictive marker of taxol efficacy as well as a new drug target for future anti-cancer therapy.

## INTRODUCTION

Taxol is a natural compound originally isolated from the Pacific yew tree *Taxus brevifolia*, and is one of the most successful chemotherapeutic drugs in the treatment of human cancer (1,2). Taxol improves overall survival rates in patients with breast, ovarian, lung cancer, and other types of solid tumor (3,4). The mechanism of action of taxol has previously been linked to mitotic arrest (5,6), apoptosis, and more recently to defects in mitotic spindle assembly and chromosome segregation in primary breast tumors (7). However, unanswered questions remain that the mechanism of action in mitotic arrest, which is ubiquitously present in all cells, may not be sufficient to explain the tumor specificity, clinical efficacy, and side effects of taxol (8–11).

We and others have previously identified a breast and ovarian cancer initiating gene *GT198* (gene symbol *PSMC3IP*, also known as Hop2 or TBPIP) (12–15). Here, we report that taxol directly interacts with GT198 and inhibits its DNA-binding activity. GT198 is a DNA-binding protein that functions as a transcriptional coactivator stimulating nuclear receptor-mediated gene activation (16,17). GT198 also acts as a crucial DNA repair factor by participating in homologous DNA recombination (12,13,18). Germline mutations in *GT198* have previously been identified in familial and early-onset breast and ovarian cancer (14,19), as well as in familial ovarian disease (20). Somatic mutations in *GT198* are prevalent and recurrent in sporadic breast, ovarian, prostate, uterine, and fallopian tube cancers, where they lead to constitutive activation of GT198 in transcription (14,15). In sporadic breast and ovarian cancers, *GT198* is mutated in tumor microenvironment. The GT198 mutant cells include pericyte stem cells and the decedent vascular smooth muscle cell lineage. These include myoepithelial cells and adipocytes in breast cancer (21), hormone-producing luteinized theca cells in ovarian cancer (22), myofibroblasts in prostate and bladder cancers (23), pericytes in skin and brain tumors as well as in multiple common solid tumors (24).

The expression of GT198 in normal tissues (16,22), is intriguingly correlated with the spectrum of taxol-induced clinical side effects (4,25). We then did a preliminary screen of commonly used chemotherapy drugs using GT198 as a protein target, and found that GT198 is inhibited by a number of oncology drugs including doxorubicin, etoposide, and taxol (23). In this study, we sought to detailed analysis of GT198 as a direct target of taxol.

## RESULTS

### Taxol binds to the DNA-binding domain of GT198

To test for a direct interaction *in vitro*, recombinant His-tagged GT198 proteins were analyzed for binding kinetics on 96-well plates using a commercially available Oregon green-labeled taxol (Figure 1a). A previously reported biotinylated taxol (26), is also analyzed for labeled taxol interaction with GT198 in cells. We first verified the biotin moiety on biotinylated taxol by streptavidin binding (Figure 1b). The integrity of both Oregon green- and biotin-labeled taxols were further evaluated by cell survival dose-response curves, which showed that both the green- and biotin-labeled taxols retain cytotoxicity albeit with much reduced efficacy (EC_50_=67.1 nM and 70.7 nM, respectively) compared to unlabeled taxol (EC_50_=3.3 nM) (Figure 1c). The addition of carbon C7 side chains reduced but did not abolish the binding of labeled taxol to cellular targets, allowing for subsequent testing of the labeled taxols. The true *in vitro* binding affinity of taxol can only be measured through competition of DNA binding to GT198 by unlabeled taxol.

**Figure 1.**
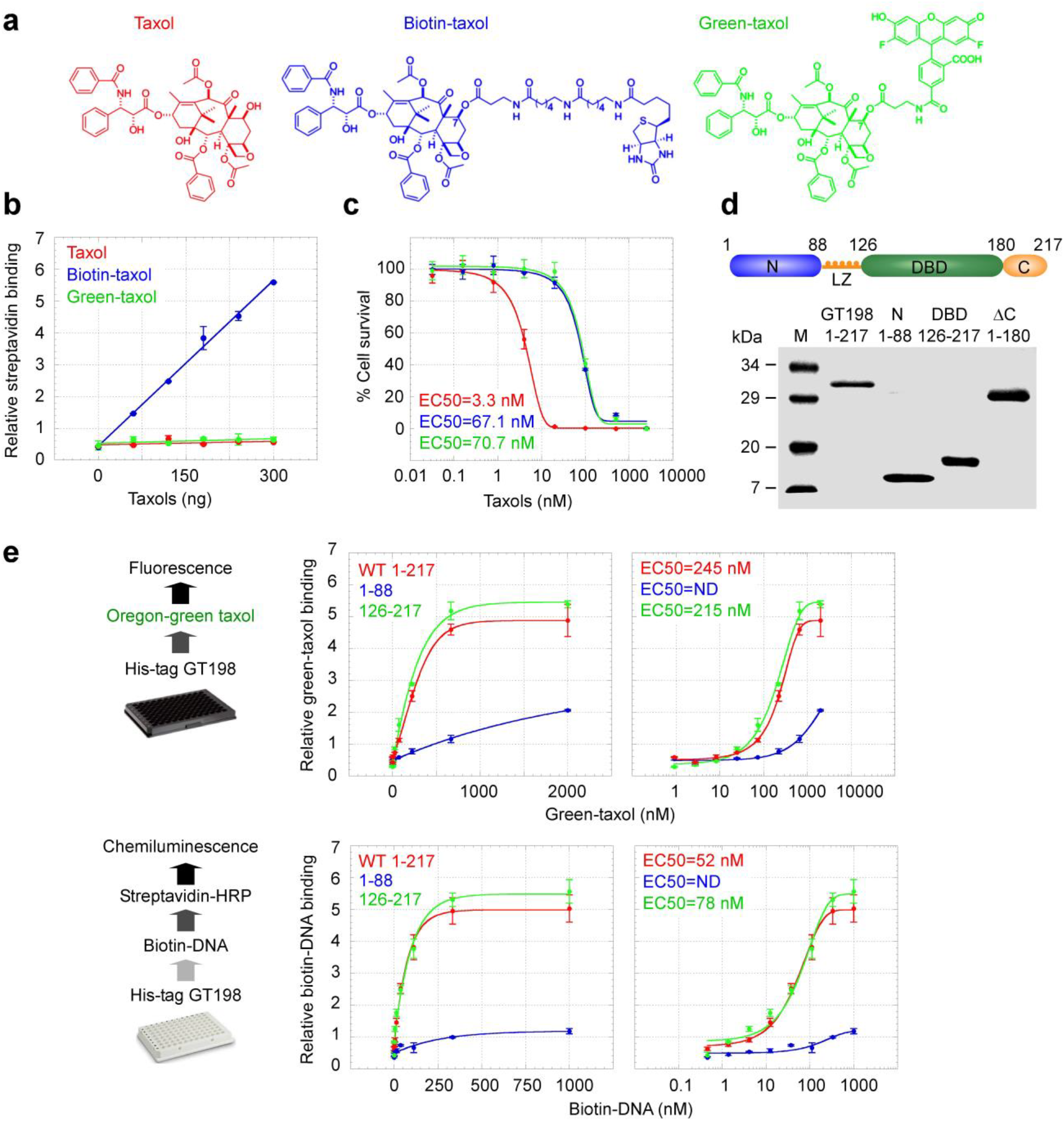
Taxol binds to the DBD of GT198. (**a**) Chemical structures of taxol (red), biotinylated taxol (blue), and Oregon green-labeled taxol (green). (**b**) Biotin was detected by streptavidin-conjugate binding to taxol at the indicated amounts. (**c**) Dose-response curves for three taxols measured by HeLa cell survival using cell viability assays. (**d**) Diagram of GT198 domain structures with purified His-tagged GT198 proteins shown below. Numbers indicate amino acids. (**e**) Top panels: *In vitro* binding of Oregon green-taxol and His-tagged GT198 proteins. Data are graphed in both linear (left) and log scales (right). EC_50_ values are the taxol concentrations for 50% binding to GT198 proteins. Bottom panels: The EC_50_ for DNA binding to GT198 proteins was measured using biotinylated DNA and detected by streptavidin-conjugate. Data represent mean ± s.e.m of duplicate experiments (n = 2).

First, we measured the binding affinity of green-taxol to His-tagged GT198 and its fragment proteins (Figure 1d), and obtained an EC_50_ of 245 nM for the wildtype GT198 (Figure 1e). The binding site was mapped using truncated GT198 protein fragments (Figure 1d). GT198 is a small homodimeric DNA-binding protein. Each monomer contains 217 amino acids including an N terminal domain, a leucine zipper (LZ) dimerization domain (16), a DNA-binding domain (DBD) capable of binding to either single- or double-stranded DNA (12,15), and a C terminal auto-regulatory domain (15). Only the GT198 DBD domain was required for the interaction of green-taxol (Figure 1e), whereas the N and C terminal domains were dispensable (Figure 1e and Supplementary Figure S1a). We also measured the DNA binding affinity of GT198 using a biotin-labeled oligonucleotide and obtained an EC_50_ value of 52 nM (Figure 1e). This high binding affinity of DNA to GT198 permitted the sensitive detection in subsequent competition assays (Figure 2).

**Figure 2.**
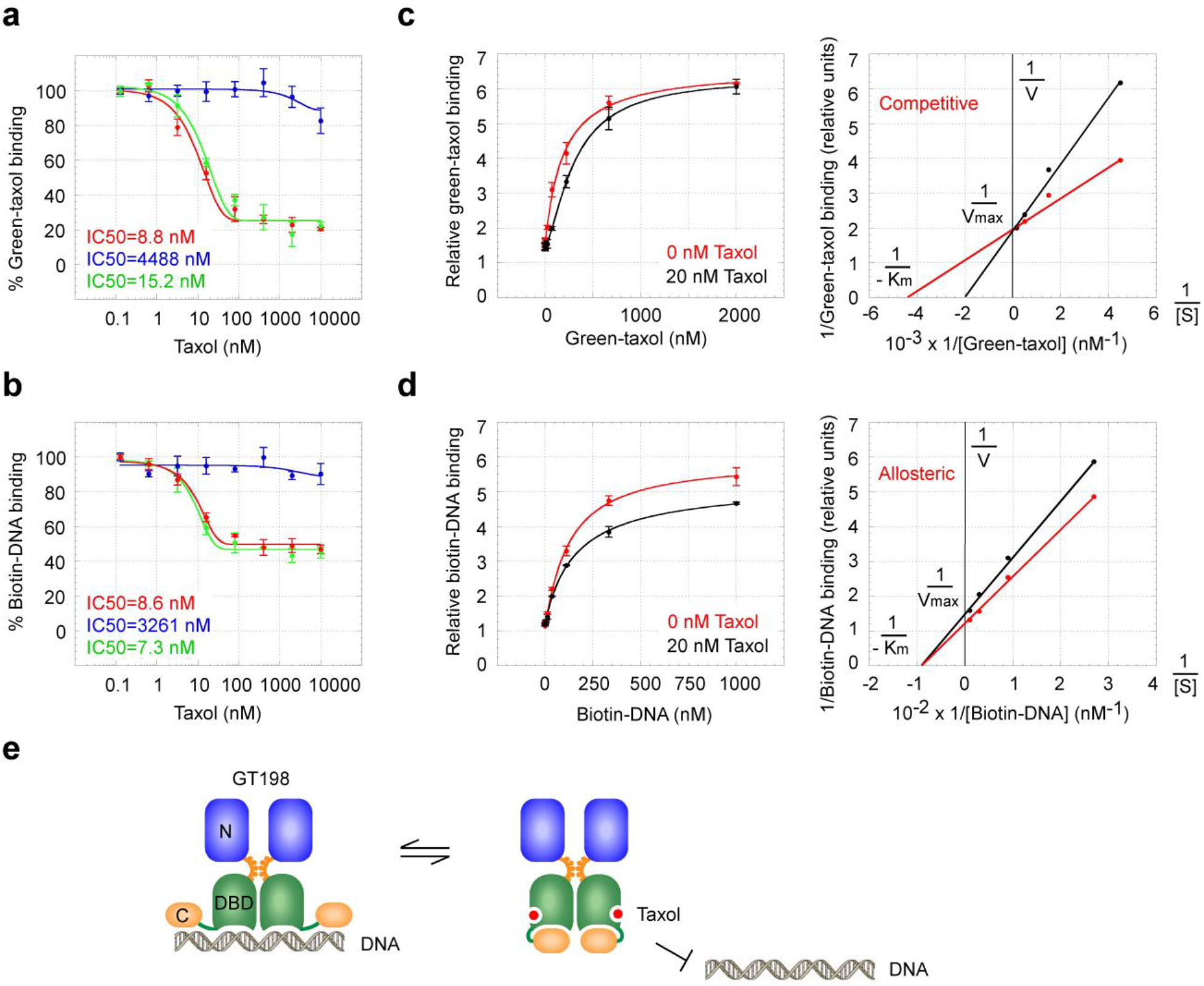
Taxol inhibits the DNA binding of GT198. (**a**) Competition binding of 400 nM green-taxol with increasing concentration of taxol to GT198 (red), its N terminal domain 1-88 (blue), and the DBD fragment 126-217 (green). IC_50_ values are the taxol concentrations for 50% inhibition of green-taxol binding. (**b**) Competition binding of 150 nM biotinylated DNA to GT198 proteins with increasing concentrations of taxol. Background binding by the N-terminal domain was not competed. (**c**) Green-taxol binding to GT198 in the presence (black) or absence (red) of 20 nM taxol. The competitor taxol shifts the curve to the right. Double reciprocal plot using the means of high concentration data points (74, 222, 666, 2000 nM) in the right panel shows competitive inhibition. (**d**) Biotinylated DNA binding to GT198 in the presence (black) or absence (red) of 20 nM taxol, showing allosteric non-competitive inhibition. Data represent mean ± s.e.m of duplicate experiments (n = 2). (**e**) Hypothetical model of taxol (red) interaction with the GT198 dimer at its DBD that induces allosteric inhibition of DNA binding.

Furthermore, we confirmed that the GT198 DBD was required for binding to biotin-labeled taxol using GST-GT198 fusion proteins (Supplementary Figure S2a-b), as well as endogenous GT198 protein immunoprecipitated from HeLa nuclear extracts (EC_50_=176 nM) (Supplementary Figure S1b). Although the binding affinities of biotin- and green-labeled taxols to GT198 may be comparable (Figure 1c), biotin detection is less sensitive than fluorescence and we were not able to assay using small quantity of His-tagged GT198 proteins, but only able to measure its interaction using GST fusions or immuneprecipitated cellular GT198. Together from the data above, we conclude that labeled taxol binds to the DBD of GT198.

### Taxol inhibits the DNA binding of GT198

Binding of unlabeled taxol to GT198 was then assessed indirectly using competition binding assays, in which increasing concentrations of unlabeled taxol compete for binding of labeled taxol. The results suggest that unlabeled taxol binds to GT198 at a much higher binding affinity since it competes for green-taxol with an IC_50_ of 8.8 nM using full-length GT198 (Figure 2a). This fold-difference in binding affinity between unlabeled and labeled taxol is comparable to the observed fold-difference in their cytotoxicities (Figure 1c). Consistently, DNA binding to GT198 was competed off using unlabeled taxol with an IC_50_ of 8.6 nM (Figure 2b), suggesting potent inhibition of GT198 DNA binding activity by unlabeled taxol. Double-reciprocal plots revealed competitive binding of GT198 between labeled and unlabeled taxols (Figure 2c). The constant V_max_ with increased K_m_ in the presence of taxol suggests that labeled green-taxol binds to the same site on GT198 as does unlabeled taxol. In contrast, inhibition of DNA binding by taxol was allosteric or non-competitive (Figure 2d). The reduced V_max_ with constant K_m_ for DNA binding in the presence of taxol suggests that taxol and DNA bind to different sites on the GT198 DBD (Figure 2d). The inhibition of DNA binding by taxol may result from a protein conformational change (**modeled in** Figure 2e). Of note, a rather incomplete inhibition of DNA binding was observed even using high concentrations of taxol (Figure 2b), further suggesting an allosteric mechanism of inhibition. Potentially, even incomplete inhibition could have sufficient impact at the functional level, since the binding affinity is rather high. Taken together, these results suggest that taxol directly inhibits the DNA binding of GT198 *in vitro* at single digit nanomolar concentrations, within a concentration range effective for cytotoxicity (Figure 1c), and is also consistent with observed drug efficacy in clinical chemotherapy.

### Taxol colocalizes with GT198 in interphase nuclei

We further examined the cellular distribution patterns of GT198 and taxol. In GT198 transfected HeLa cells, biotinylated taxol colocalizes with overexpressed GT198 and its DBD fragment but not with its N terminal domain (Figure 3a). As previously reported, GT198 is nuclear and its fragments are cytoplasmic (15). The colocalization is consistent with the observed binding of taxol to the DBD of GT198. The punctate pattern of endogenous GT198 coincided more precisely than overexpressed GT198 to the biotin-labeled taxol staining in interphase nuclei (Figure 3b). In mitotic phase cells, endogenous GT198 and taxol staining were significantly increased relative to the interphase suggesting that the GT198 expression is correlated with the cell cycle (Figure 3b). Similarly, green-taxol and GT198 nuclear staining patterns closely coincided in interphase nuclei, although additional cytoplasmic staining was observed with green-taxol (Figure 3c). This cytoplasmic signal was not due to microtubule binding as detected using anti-α-tubulin, since α-tubulin staining did not coincide with green-taxol in either interphase or mitotic phase cells (Figure 3c). Notably, the nuclear but not the cytoplasmic staining of green-taxol was blocked by unlabeled taxol (Supplementary Figure S3), suggesting this cytoplasmic signal is perhaps non-specific. Thus, taxol colocalizes specifically with GT198 but not microtubules in HeLa cells.

**Figure 3.**
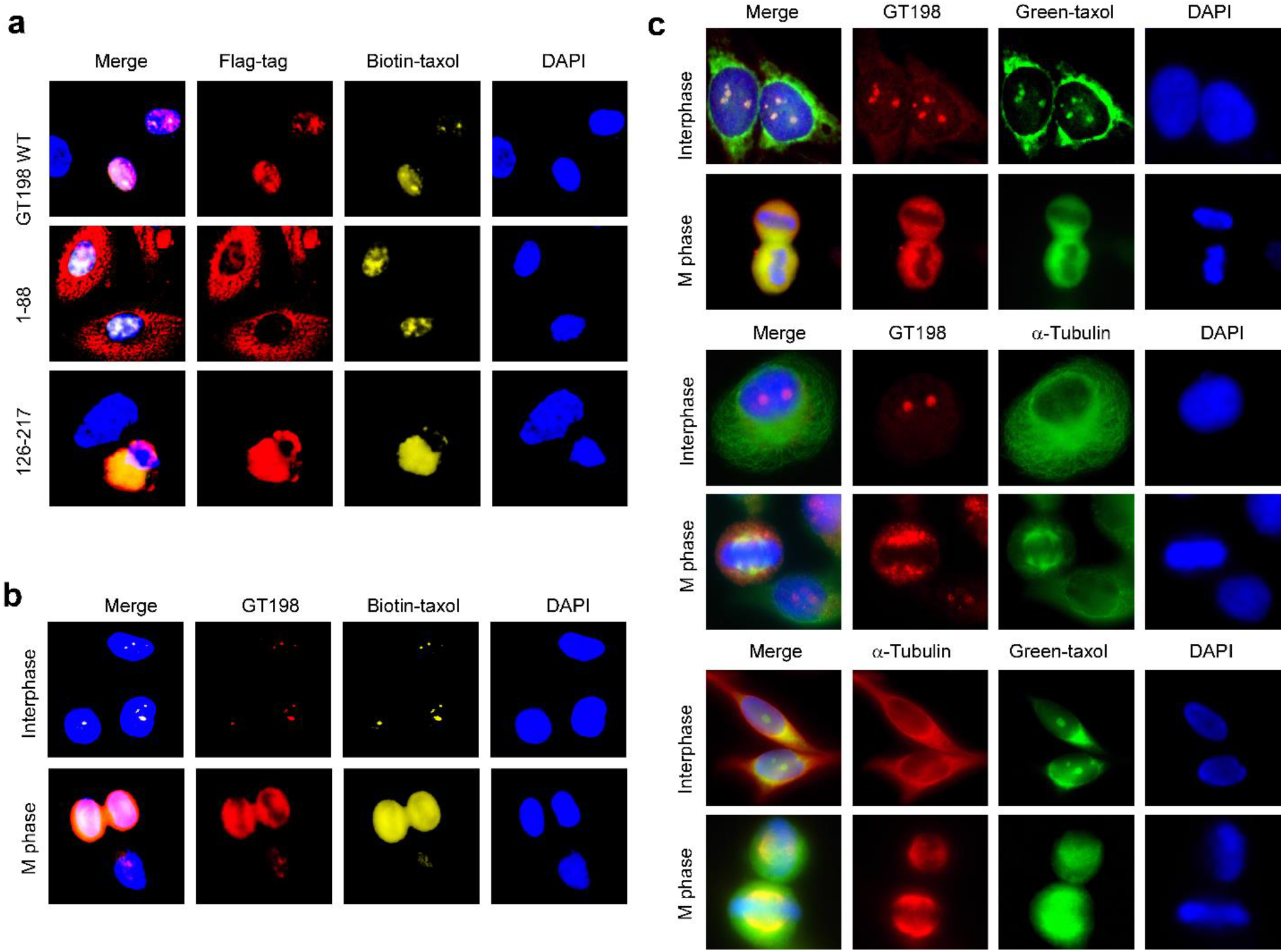
Taxol and GT198 colocalize in HeLa cells. (**a**) Fluorescent double staining of HeLa cells in which transfected GT198 or its fragment proteins (0.1 μg plasmid DNA transfected) were detected by anti-Flag antibody (1:1000) in red. Wildtype GT198 and the DBD are expressed in nuclei and the N-terminus is in cytoplasm as previously reported (15). The staining of biotinylated taxol (5 μM) was detected by streptavidin-PE in yellow. (**b**) Endogenous GT198 in interphase or mitotic phase was detected by affinity purified anti-GT198 antibody (1:150) in red, co-stained with biotinylated taxol (5 μM), and streptavidin-PE in yellow. (**c**) Green-taxol (1 μM) in green, endogenous GT198 detected by anti-GT198 (1:150) in red, and anti-α-tubulin (1:150) in red or green, were co-stained in combinations to compare localization in interphase and mitotic phase of untransfected HeLa cells. An extended view of GT198 and green-taxol is shown in the supplementary materials (Supplementary Fig. 3). Secondary antibodies were Alexa 595 (red) and Alexa 488 (green)-conjugated anti-mouse or anti-rabbit antibodies. Cells were counterstained with DAPI in blue.

### Decreased GT198 desensitizes Taxol-induced cell death and taxol inhibits GT198 foci formation

To analyze the functional consequence of taxol binding to GT198, we reduced GT198 expression using transient transfection of siRNA in HeLa cells. GT198 siRNA desensitized, whereas GT198 overexpression sensitized cells to the cytotoxic effects of taxol (Figure 4a). Similarly, in mouse P19 stem cells, the stably expressed DBD fragment of GT198 (aa 126-217) desensitized cytotoxicity of taxol relative to stable expression of GT198 (Figure 4b), consistent with the notion that DBD acts as a dominant negative of wild type GT198 (15). In particular, a significant portion of DBD-transfected stable cells remained viable even after exposure to high concentrations of taxol (Figure 4b), suggesting that reduced GT198 activity may render taxol less effective. GT198 is also a DNA repair factor, and forms nuclear foci upon γ-irradiation (Figure 4c). Strikingly, taxol treatment results in GT198 cytoplasmic translocation and disrupts GT198 foci formation in irradiated cells (Figure 4c). This finding is consistent with the observation that translocation of DNA repair proteins to the cytoplasm occurs following taxol treatment (10). Since GT198 cytoplasmic translocation is characteristic of wild type GT198 inhibition (15,22), this result suggests that taxol inhibits GT198 activity, including its function in DNA repair.

**Figure 4.**
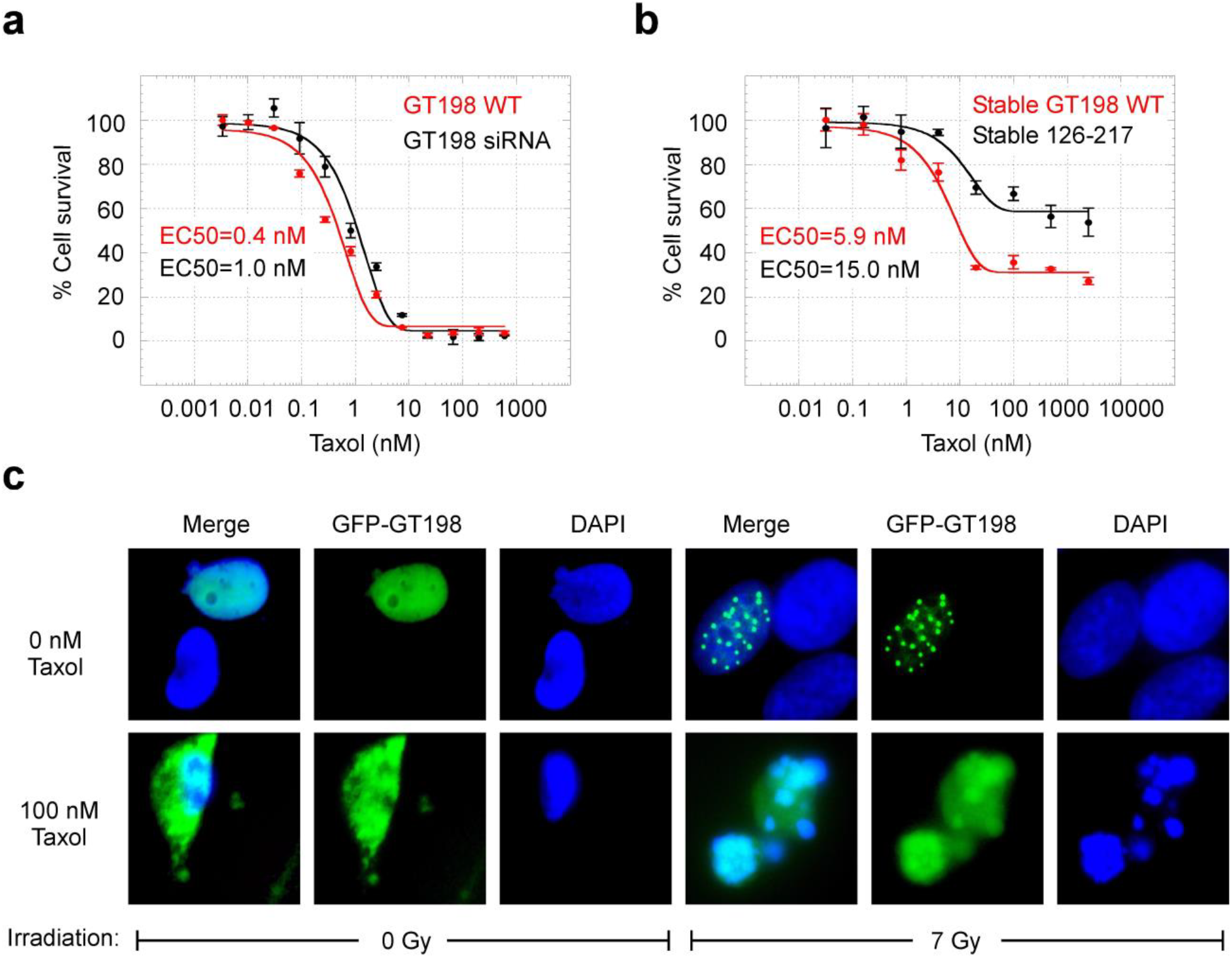
GT198 level is correlated with taxol cytotoxicity and taxol inhibits GT198 nuclear foci formation. (**a**) HeLa cells were transfected with 0.4 μg/well plasmid of GT198 (red) or 100 nM GT198 siRNA (black), and treated with indicated concentrations of taxol for 72 hours before analysis using cell viability assay. (n = 2), *P* value=0.0449. (**b**) Mouse P19 stem cells were stably transfected with GT198 (red) or the DBD fragment 126-217 (black) and treated with taxol for 72 hours before measuring cell viability. (n = 2), *P* value=0.0066. Data represent mean ± s.e.m of duplicate experiments. (**c**) HeLa cells were transfected with GFP-GT198 (green), and then treated with 100 nM taxol for 24 hours. GFP-GT198 foci formation was induced by γ-irradiation at 7 Gy followed by recovery for 4 hours before imaging. Cells were counterstained with DAPI.

## DISCUSSION

In summary, our results provide the first evidence that a DNA repair factor and oncoprotein GT198 is a direct target of taxol. From a clinical perspective, the study provides anticipated rationale for the clinical side effects of taxol. In human cancer patients, taxol-induced lymphocyte toxicity (27), neutropenia (4), testicular shrinkage (28), and ovarian damage (29), are potentially correlated with the endogenous GT198 protein expression specifically in testis, ovary, spleen, and thymus (16,22). The acute taxol toxicity in rodents was also found in ovary (30), testis, hematopoietic, and lymphoid systems (31). In contrast, the low toxicity of taxol in major organs is correlated with undetectable GT198 protein level therein (16). Moreover, GT198 stimulates glucocorticoid receptor-mediated transcription (16). The glucocorticoid receptor ligand dexamethasone is an effective premedication prior to taxol (25), potentially due to a compensation of taxol-induced GT198 loss in normal cells.

In human ovarian cancer, for example, GT198 is overexpressed in the luteinized theca cells and causes steroid hormone overproduction in the tumor microenvironment (22). Mutant luteinized theca cells provide hormone stimuli to recapitulate epithelial tumor cells. GT198 inhibition by taxol would be predicted to suppress steroid hormone for cancer initiation thereby result in high clinical efficacy. Consistently, in human breast cancer, GT198 is expressed in mutant pericyte stem cells, the decedent fibroblasts, adipocytes, and myoepithelial cells (21). Taxol treatment of human breast cancer would inhibit the cancer-inducing cell lineages in tumor microenvironment and lead to observed clinical efficacy in the treatment. In addition, GT198 regulates androgen receptor-mediated transcription (16), and taxane chemotherapy that would suppress GT198 has already been shown to impair androgen receptor-mediated transcription in prostate cancer (9,32). At the cellular level, taxol inhibits microtubule trafficking of a number of DNA repair proteins (10). Inhibited GT198 within the DNA repair complex could potentially lead to observed defects in microtubule trafficking (33). Finally, GT198 is highly expressed in cultured cancer cell lines (Supplementary Figure S4) (16), and in xenografted mouse tumor (24), which could underlie the historic success in taxol identification using GT198-positive cell lines or xenografted tumors (1,34). Together these observations strongly support that GT198 is a direct protein target of taxol. In the future, GT198 may be a useful prognostic marker in predicting the clinical efficacy of taxol or taxane in human cancer treatment. GT198 can also serve as a therapeutic drug target to identify new drugs.

## ACKNOWLEDGEMENTS

We thank Dr. Susan B. Horwitz for critical comments on this work. We thank Dr. Nita J. Maihle for manuscript discussion. We thank Drs. Mingqiang Ren, Xiongjie Jin, and Hasan Korkaya for reagents and help. This work was supported in part by the Georgia Cancer Coalition Distinguished Cancer Scholar Award to L.K., the National Institutes of Health grant CA062130 to N.F.M. and GM094313 to M.L.G.

## SUPPLEMENTARY INFORMATION

This article has a supplementary information in a separate PDF file.

---

**Supplementary Figure S1.**
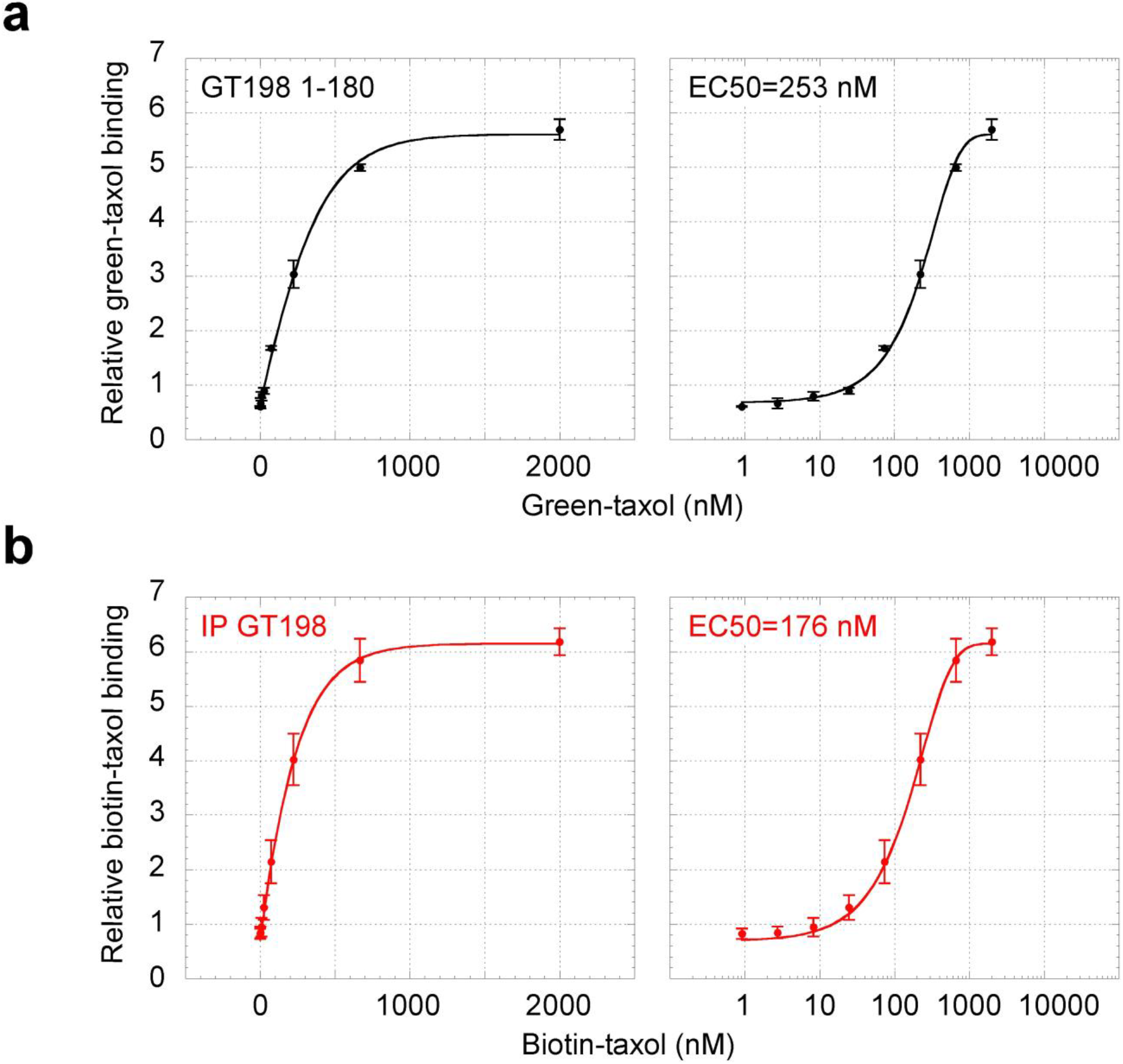
Taxol binds to GT198. **(a)** *In vitro* binding of green-taxol and His-tagged GT198 1-180 lacking the C terminal domain. The C-terminus is thus not required for taxol interaction. Dose-response curves are graphed in linear (left) and log scales (right). **(b)** Dose-response curves of *in vitro* binding of biotinylated taxol and GT198 immunoprecipitated from HeLa nuclear extracts. Biotinylated taxol was detected by streptavidin-conjugated HRP. Data represent mean ± s.e.m of duplicate experiments (n = 2).

**Supplementary Figure S2.**
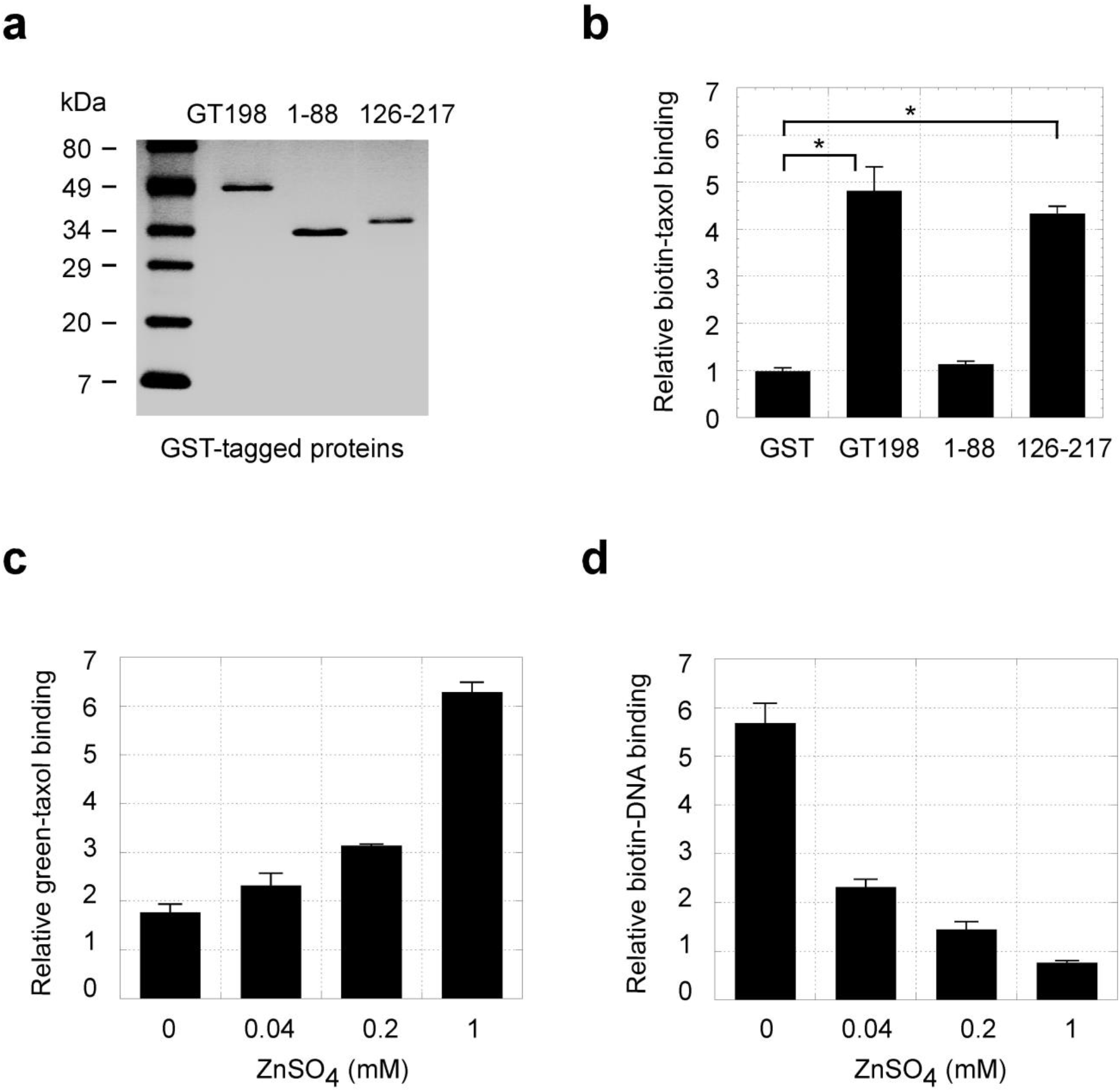
Taxol binds to GST-GT198 and the effect of zinc concentrations on GT198 binding. **(a)** Coomassie blue staining of purified GST-tagged GT198 and its protein fragments. **(b)** GST pull down in the presence of biotinylated taxol (2 mM) shows that taxol interacts with DBD-containing GT198 proteins. (n = 2), *P* * < 0.05. **(c)** The binding of His-tagged GT198 and green-taxol (400 nM) is zinc-dependent. (n = 2), *P* value=0.0061. **(d)** Binding of GT198 by biotinylated DNA (150 nM) is inhibited by zinc. (n = 2), *P* value=0.0079. To achieve maximal green-taxol binding, the zinc concentration is optimized to test green-taxol effect on GT198 with DNA interaction (see Methods). The presence of potential zinc fingers in GT198 remains to be determined.

**Supplementary Figure S3.**
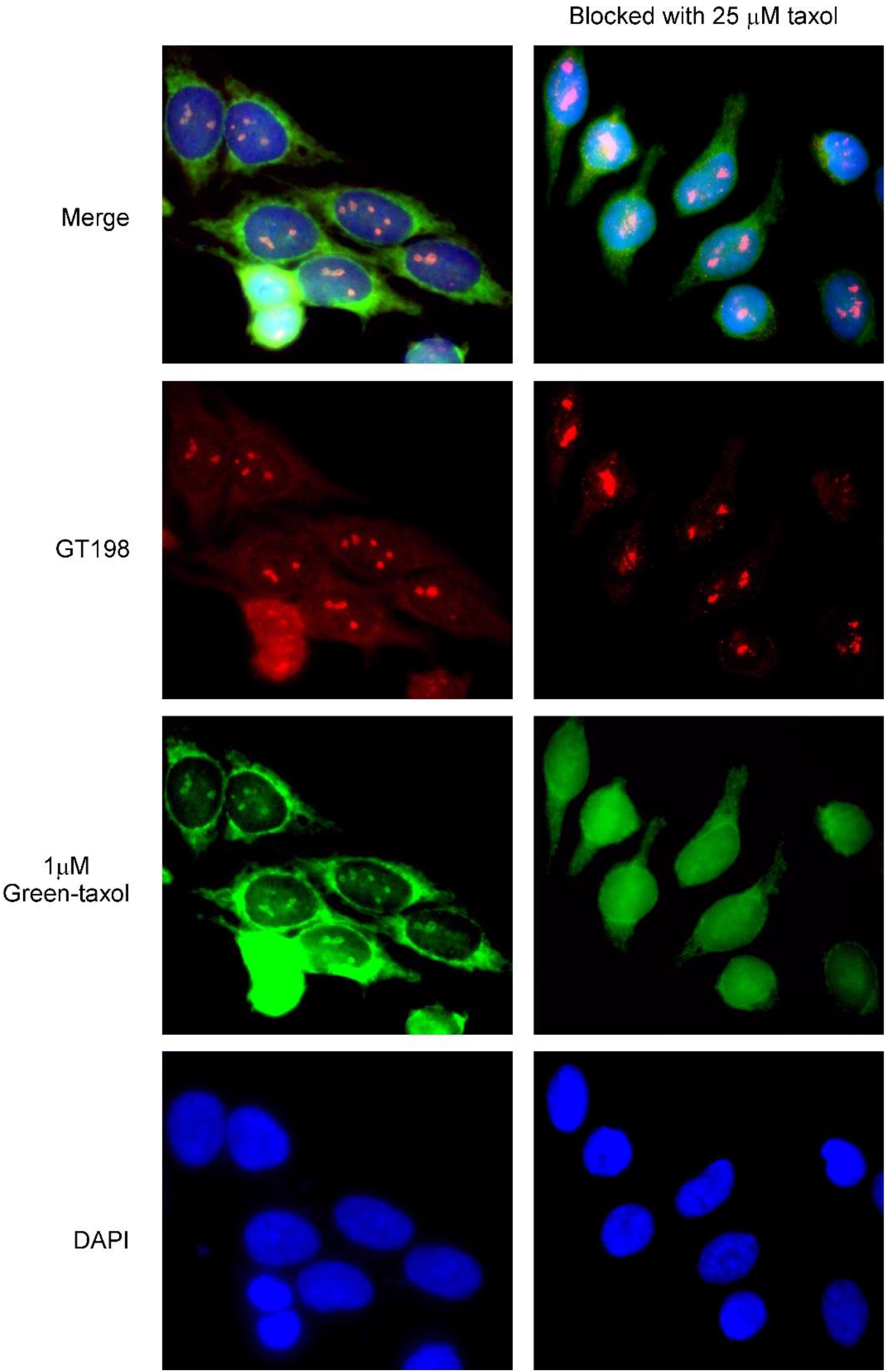
Green-taxol staining is specific to nuclear GT198 in HeLa cells. The left panels show an extended view of an image from Figure 3c, in which HeLa cells were stained with green-taxol (1 μM) in green, anti-GT198 (1:150) in red, and DAPI in blue. Two newly divided cells in lower left corner have stronger staining of GT198 as well as green-taxol. The right panels show staining under the same conditions except with additional 25 μM unlabeled taxol to block green-taxol. Nuclear but not cytoplasmic non-specific staining of green-taxol is blocked.

**Supplementary Figure S4.**
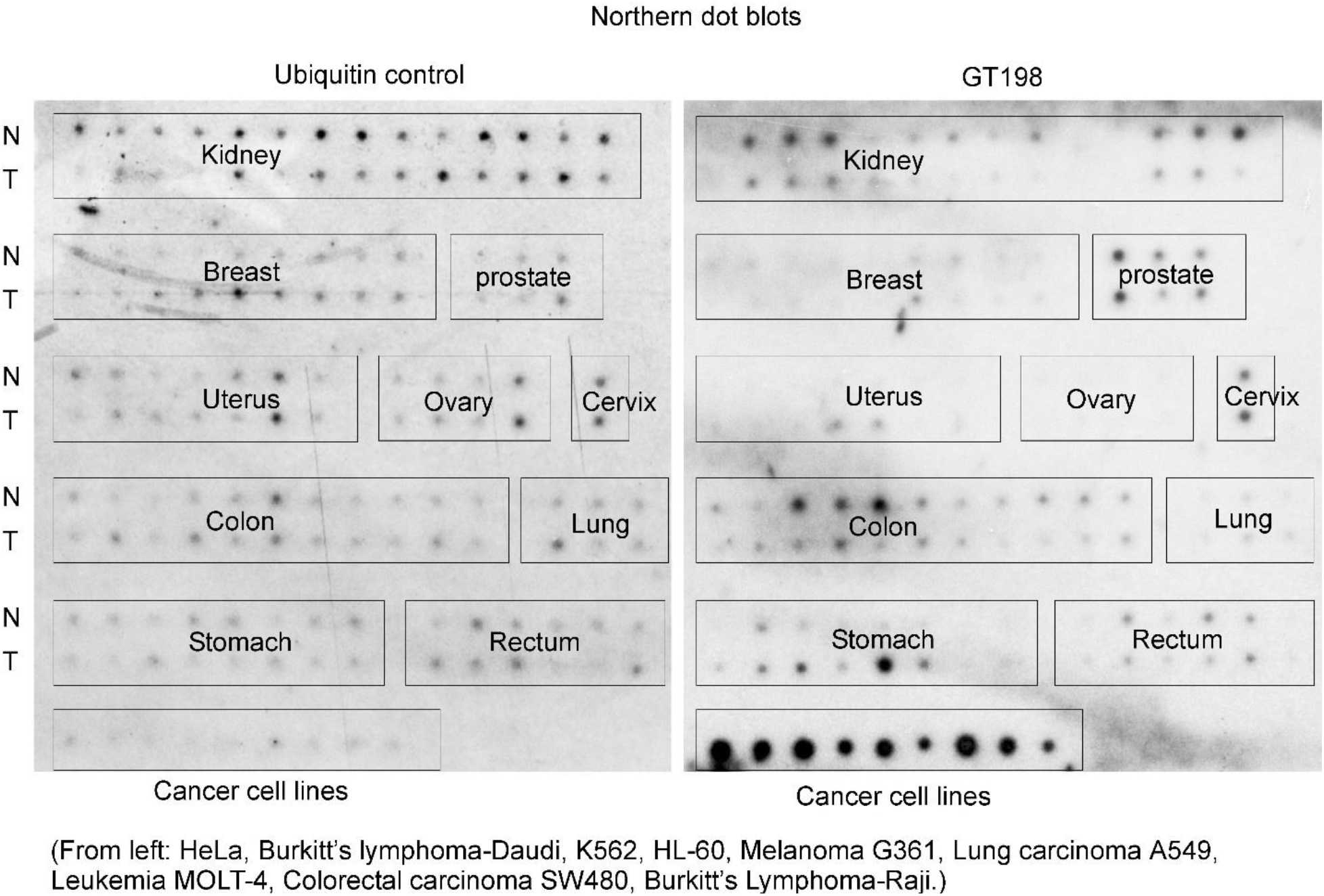
GT198 mRNA expression in human solid cancers. Northern dot blot analyses of GT198 mRNA expression in paired human tumors (T), adjacent normal tissues which may contain altered tumor microenvironment (N), and in selected cancer cell lines (Clontech, 7840-1). Ubiquitin in the left panel serves as a control. GT198 expression is at low levels in most primary tumors but is significantly higher in cancer cell lines.

## METHODS

### GT198 protein purification

N-terminal His-tagged recombinant human GT198 proteins including wild type full length GT198 (aa 1-217), N terminus (aa 1-88), DBD fragment (aa 126-217), and C-terminus deletion (aa 1-180) were expressed in *E. coli* BL21(DE3)pLysS and purified through Ni-NTA-agarose (Qiagen, #30210). Proteins were eluted by 200 mM imidazole, desalted and concentrated using Amicon YM-10 spin columns before use. The glutathione *S*-transferase (GST) fusion GT198 proteins were expressed in the same *E. coli* strain and purified using Glutathione Sepharose columns (Amersham Pharmacia, #17-0756-01). GST proteins were retained on the Sepharose beads in binding assays. For immunoprecipitated GT198, 5 μl anti-GT198 rabbit serum were captured by 10 μl Protein A/G agarose (Santa Cruz, sc-2003), washed, and incubated at 4°C for overnight with 200 μl HeLa (ATCC, CCL-2) nuclear extracts in a total volume of 1.2 ml buffer. The immune complex was washed and subjected to binding assays.

### Green-taxol binding and competition assays

The binding of Oregon green-labeled taxol (Molecular Probe, Oregon Green^®^ 488 Taxol, P-22310) to GT198 was detected through analyzing green fluorescence. Black 96-well plates (Thermo Scientific, #7805) were coated to dry overnight at 37°C with 400 ng/well of recombinant His-tagged GT198 proteins together with 5 μg/well of purified BSA (NEB) in a volume of 50 μl. BSA alone was included as a control for background. No subtraction of background was needed due to very low background. Duplicate wells were used for each experimental point (n = 2). The GT198-coated plates were blocked with 5% BSA in PBS with 0.1% Triton X-100 (TPBS) for 1 h. The binding was carried out in 100 μl/well for 4 hrs to overnight at 4°C using serial diluted green-taxol (0.91, 2.74, 8.23, 24.69, 74.07, 222.22, 666.66, 2000 nM) in the binding buffer (20 mM TrisHCl, pH 7.5, 50 mM NaCl, 75 mM KCl, 0.5 mM MgCl_2_, 0.05% Triton X-100, 10% glycerol, 1 mM dithiothreitol) containing 1 mM ZnSO_4_. For competition assays, binding was carried out with 400 nM green-taxol and serial diluted unlabeled taxol (0.128, 0.64, 3.2, 16, 80, 400, 2000, 10000 nM) (Novation). The plates were then washed five times with TPBS within 20 min. Green fluorescence was measured under the wavelengths of 496 nm excitation and 524 nm emission in a Tecan Safire microplate reader. See the Supplementary Information page for further details.

### DNA binding and competition assays

The binding of biotinylated DNA to GT198 was detected by chemiluminescence. The DNA binding to GT198 is non-sequence specific (15). A single-stranded 25-mer biotin-oligonucleotide [Biotin]-cctggggttgctgaggtcctggcag was used in the assay since it is sufficient to bind one GT198 dimer. White MicroLite™ 2+ 96-well plates (Thermo Scientific, #7572) were similarly coated with His-tagged GT198 proteins and blocked with BSA as above. The binding in duplicates (n = 2) was carried out for 4 hrs to overnight at 4°C using serial diluted Biotin-DNA (0.45, 1.3, 4.1, 12.3, 37.0, 111.1, 333.3, 1000 nM) in the binding buffer above without ZnSO_4_. For competition assays, binding was carried out with 150 nM Biotin-DNA and serial diluted taxol (0.128, 0.64, 3.2, 16, 80, 400, 2000, 10000 nM) in the binding buffer containing 0.1 mM ZnSO_4_. After binding, the plates were washed three times with TPBS in 30 min. The plates were then incubated with streptavidin-conjugated horseradish peroxidase (HRP) (Roche Molecular Biochemicals, #1089153) at 1 U/ml for 1 h at 4°C and further washed three times with TPBS in 30 min. Bound DNA were detected by chemiluminescence with 50 μl/well ECL detection reagents (Amersham Pharmacia Biotech) in a Dynex MLX luminometer.

### Binding assays using GST-GT198 and immunoprecipitated GT198

*In vitro* binding assays using GST fusions were performed in duplicates by incubating GST-GT198 resins (10 μl, 2 μg) and biotinylated taxol (2 μM) in the binding buffer containing 1 mM ZnSO_4_ for overnight at 4°C. For immunoprecipitated GT198, experiments in duplicates were performed by incubating agarose beads-captured immune complex and biotinylated taxol (2 μM) in the binding buffer for overnight at 4°C. Washed beads were incubated with streptavidin-peroxidase (1 U/ml) for 1 h at 4°C and further washed three times before transfer to 96-well plates to measure. Bound taxol was detected by chemiluminescence with 50 μl/well ECL detection reagents in a Dynex MLX luminometer.

### Kinetic data analysis

The binding EC_50_ values are the 50% of maximal binding concentrations of green-taxol or DNA. The inhibition IC_50_ values are the taxol concentrations of 50% inhibition of green-taxol or DNA binding. In cell viability assays, EC_50_ values are the taxol concentrations that reduce total cell viability by 50%. The EC_50_ and IC_50_ values were calculated through Kaleidagraph software (Synergy Software) using nonlinear regression sigmoidal dose-response curve fit. The equation for calculation is y = m1 + (m2 – m1)/(1 + (x /m3)^m4^), where m1 is the minimum, m2 is the maximum, m3 is the EC_50_ or IC_50_ value calculated by Kaleidagraph, and m4 is the slope at midpoint of the curve. However, data points were drawn with better fitted slopes using nonlinear regression sigmoidal curve fit provided by Kaleidagraph software: y = m1 + m2/(1 + exp(-(x-m3)/m4)), where exp = е^x^, although this equation was less tolerant in calculation and was not used to calculate values. The analysis of V_max_ and K_m_ in competitive or non-competitive binding assays was carried out in double-reciprocal plots, using means of four high concentration data points, omitting points at lower concentrations with reciprocal values off-scale.

### Cell viability assays

HeLa cells (ATCC, CCL-2) were maintained in DMEM supplemented with 10% fetal bovine serum, 100 U/ml penicillin and 0.1 µg/µl streptomycin. Cells were incubated in 5% CO_2_ at 37°C. For transient transfections, GT198 plasmid (0.4 μg/well) or GT198 siRNA (100 nM) was transfected into HeLa cells in 48-well plates for 4 hrs before the cell viability assay. Mouse embryonal carcinoma P19 cells (ATCC, CRL-1825) were maintained in α-MEM supplemented with 2.5% fetal bovine and 7.5% bovine calf serum, and antibiotics above. Stably transfected P19 cells expressing GT198 and its DBD-containing fragment 126-217 were selected using 400 g/ml of G418 for three weeks. Positive clones were confirmed by PCR and Western blot analyses. The cell viability assays were carried out in duplicate wells for each condition in 48-well plates after taxol treatment for 72 hrs at various concentrations (0.032, 0.16, 0.8, 4, 20, 100, 500, 2500 nM). The tetrazolium compound MTS reagent can be reduced by cells into a colored formazan product (CellTiter 96® AQueous One Solution Cell Proliferation Assay, Promega, G3580). The MTS reagent was diluted in phenol red-free medium to minimize the background and incubated with cells at 200 µl/well for 1 h at 37°C. The reacted colored medium was transferred to flat-bottom clear 96-well plates and the absorbance at 490 nm was determined using a Tecan Safire microplate reader. *P* values were determined by two-tailed paired t test.

### Immunofluorescence

Immunofluorescence double staining was carried out using 5 μM biotinylated taxol (26), or 1 μM Oregon green-labeled taxol (Molecular Probe, P-22310), together with rabbit or mouse antibodies. Secondary antibodies are Alexa 488 (green)- and Alexa 594 (red)-conjugated anti-rabbit or anti-mouse antibodies (Invitrogen, A11001, A11037, A11005). Polyclonal anti-GT198 antibody (1:150) was previously prepared from rabbits (Covance), and affinity purified using Affi-gel 10 (Bio-Rad, 153-6099). Commercial available antibodies are mouse anti-Flag (1:1000, Sigma, M2, F3165), and mouse anti-α-tubulin (1:150, Santa Cruz, sc-5286). Biotinylated taxol staining was detected by streptavidin-R-Phycoerythrin (PE) (1:1000, Invitrogen, S-866) in yellow. Flag-tagged GT198 plasmids (0.1 μg)-transfected or non-transfected HeLa cells in chamber slides were methanol fixed, blocked with 4% horse serum, and stained by primary antibody together with labeled taxol in 1% horse serum in TPBS overnight at 4°C. After washing, slides were incubated with secondary antibody or streptavidin-PE in 1% horse serum in TPBS for 1 h in the dark. Slides were counterstained with DAPI before visualization by fluorescence microscopy. The plasmid GFP-GT198 contained full-length human GT198 in a pEGFP-C3 vector (Clontech, 6082-1). HeLa cells transfected with GFP-GT198 (green) were γ-irradiated at 7 Gys and recovered for 4 hours before methanol-fixed for immunofluorescence.

### Northern dot blot analysis

The Northern dot blots containing cDNAs from primary human tumors (T) with paired adjacent normal tissues (N) and cancer cell lines were obtained from Clontech (Cancer Profiling Array, #7840-1). The blots contained normalized cDNA isolated from tumors and the corresponding adjacent normal tissues from individual cancer patients. However, adjacent normal tissues often possess early molecular changes undetectable by pathology standards. The GT198 probe was prepared by random-primed ^32^P-DNA synthesis using human GT198 full-length cDNA as template. Northern hybridization was performed according to manufacturer’s protocol before visualized by autoradiography.

### Statistics

Statistical analyses were performed using Prism software. In cell viability assays, statistical significance of differences between groups was determined by two-tailed paired t test. In GST binding assays, *P* values were determined by two-tailed unpaired t test. In binding assays with zinc, *P* values were determined by linear regression. Each binding assay was repeated for at least three times in the laboratory. Data represent mean ± s.e.m of duplicate experiments (n = 2) as indicated. Duplicates rather than triplicates were used in each experimental data point due to highly consistent *in vitro* experimental conditions that yield variations are sufficiently small.

## REFERENCES

1. Wall, M. E., and Wani, M. C. (1995) Camptothecin and taxol: discovery to clinic--thirteenth Bruce F. Cain Memorial Award Lecture. Cancer Res 55, 753–760

2. Wani, M. C., and Horwitz, S. B. (2014) Nature as a remarkable chemist: a personal story of the discovery and development of Taxol. Anticancer Drugs 25, 482–487

3. Von Hoff, D. D., Ervin, T., Arena, F. P., Chiorean, E. G., Infante, J., Moore, M., Seay, T., Tjulandin, S. A., Ma, W. W., Saleh, M. N., Harris, M., Reni, M., Dowden, S., Laheru, D., Bahary, N., Ramanathan, R. K., Tabernero, J., Hidalgo, M., Goldstein, D., Van Cutsem, E., Wei, X., Iglesias, J., and Renschler, M. F. (2013) Increased survival in pancreatic cancer with nab-paclitaxel plus gemcitabine. N Engl J Med 369, 1691–1703

4. Verweij, J., Clavel, M., and Chevalier, B. (1994) Paclitaxel (Taxol) and docetaxel (Taxotere): not simply two of a kind. Ann Oncol 5, 495–505

5. Schiff, P. B., Fant, J., and Horwitz, S. B. (1979) Promotion of microtubule assembly in vitro by taxol. Nature 277, 665–667

6. Abal, M., Souto, A. A., Amat-Guerri, F., Acuna, A. U., Andreu, J. M., and Barasoain, I. (2001) Centrosome and spindle pole microtubules are main targets of a fluorescent taxoid inducing cell death. Cell Motil Cytoskeleton 49, 1–15

7. Zasadil, L. M., Andersen, K. A., Yeum, D., Rocque, G. B., Wilke, L. G., Tevaarwerk, A. J., Raines, R. T., Burkard, M. E., and Weaver, B. A. (2014) Cytotoxicity of paclitaxel in breast cancer is due to chromosome missegregation on multipolar spindles. Sci Transl Med 6, 229ra243

8. Komlodi-Pasztor, E., Sackett, D. L., and Fojo, A. T. (2012) Inhibitors targeting mitosis: tales of how great drugs against a promising target were brought down by a flawed rationale. Clin Cancer Res 18, 51–63

9. Thadani-Mulero, M., Nanus, D. M., and Giannakakou, P. (2012) Androgen receptor on the move: boarding the microtubule expressway to the nucleus. Cancer Res 72, 4611–4615

10. Poruchynsky, M. S., Komlodi-Pasztor, E., Trostel, S., Wilkerson, J., Regairaz, M., Pommier, Y., Zhang, X., Kumar Maity, T., Robey, R., Burotto, M., Sackett, D., Guha, U., and Fojo, A. T. (2015) Microtubule-targeting agents augment the toxicity of DNA-damaging agents by disrupting intracellular trafficking of DNA repair proteins. Proc Natl Acad Sci U S A 112, 1571–1576

11. Weaver, B. A. (2014) How Taxol/paclitaxel kills cancer cells. Mol Biol Cell 25, 2677–2681

12. Enomoto, R., Kinebuchi, T., Sato, M., Yagi, H., Shibata, T., Kurumizaka, H., and Yokoyama, S. (2004) Positive role of the mammalian TBPIP/HOP2 protein in DMC1-mediated homologous pairing. J Biol Chem 279, 35263–35272

13. Petukhova, G. V., Pezza, R. J., Vanevski, F., Ploquin, M., Masson, J. Y., and Camerini-Otero, R. D. (2005) The Hop2 and Mnd1 proteins act in concert with Rad51 and Dmc1 in meiotic recombination. Nat Struct Mol Biol 12, 449–453

14. Peng, M., Bakker, J. L., DiCioccio, R. A., Gille, J. J. P., Zhao, H., Odunsi, K., Sucheston, L., Jaafar, L., Mivechi, N. F., Waisfisz, Q., and Ko, L. (2013) Inactivating mutations in GT198 in familial and early-onset breast and ovarian cancers. Genes Cancer 4, 15–25

15. Peng, M., Yang, Z., Zhang, H., Jaafar, L., Wang, G., Liu, M., Flores-Rozas, H., Xu, J., Mivechi, N. F., and Ko, L. (2013) GT198 splice variants display dominant negative activities and are induced by inactivating mutations. Genes Cancer 4, 26–38

16. Ko, L., Cardona, G. R., Henrion-Caude, A., and Chin, W. W. (2002) Identification and characterization of a tissue-specific coactivator, GT198, that interacts with the DNA-binding domains of nuclear receptors. Mol Cell Biol 22, 357–369

17. Satoh, T., Ishizuka, T., Tomaru, T., Yoshino, S., Nakajima, Y., Hashimoto, K., Shibusawa, N., Monden, T., Yamada, M., and Mori, M. (2009) Tat-binding protein-1 (TBP-1), an ATPase of 19S regulatory particles of the 26S proteasome, enhances androgen receptor function in cooperation with TBP-1-interacting protein/Hop2. Endocrinology 150, 3283–3290

18. Cho, N. W., Dilley, R. L., Lampson, M. A., and Greenberg, R. A. (2014) Interchromosomal homology searches drive directional ALT telomere movement and synapsis. Cell 159, 108–121

19. Schubert, S., Ripperger, T., Rood, M., Petkidis, A., Hofmann, W., Frye-Boukhriss, H., Tauscher, M., Auber, B., Hille-Betz, U., Illig, T., Schlegelberger, B., and Steinemann, D. (2017) GT198 (PSMC3IP) germline variants in early-onset breast cancer patients from hereditary breast and ovarian cancer families. Genes Cancer 8, 472–483

20. Zangen, D., Kaufman, Y., Zeligson, S., Perlberg, S., Fridman, H., Kanaan, M., Abdulhadi-Atwan, M., Abu Libdeh, A., Gussow, A., Kisslov, I., Carmel, L., Renbaum, P., and Levy-Lahad, E. (2011) XX ovarian dysgenesis is caused by a PSMC3IP/HOP2 mutation that abolishes coactivation of estrogen-driven transcription. Am J Hum Genet 89, 572–579

21. Yang, Z., Peng, M., Cheng, L., Jones, K., Maihle, N. J., Mivechi, N. F., and Ko, L. (2016) GT198 Expression Defines Mutant Tumor Stroma in Human Breast Cancer. Am J Pathol 186, 1340–1350

22. Peng, M., Zhang, H., Jaafar, L., Risinger, J. I., Huang, S., Mivechi, N. F., and Ko, L. (2013) Human ovarian cancer stroma contains luteinized theca cells harboring tumor suppressor gene GT198 mutations. J Biol Chem 288, 33387–33397

23. Ko, L. (2019) Human solid cancer decoded. Zenodo preprint, http://doi.org/10.5281/zenodo.3236836

24. Zhang, L., Wang, Y., Rashid, M. H., Liu, M., Angara, K., Mivechi, N. F., Maihle, N. J., Arbab, A. S., and Ko, L. (2017) Malignant pericytes expressing GT198 give rise to tumor cells through angiogenesis. Oncotarget 8, 51591–51607

25. Peereboom, D. M., Donehower, R. C., Eisenhauer, E. A., McGuire, W. P., Onetto, N., Hubbard, J. L., Piccart, M., Gianni, L., and Rowinsky, E. K. (1993) Successful re-treatment with taxol after major hypersensitivity reactions. J Clin Oncol 11, 885–890

26. Lis, L. G., Smart, M. A., Luchniak, A., Gupta, M. L., Jr., and Gurvich, V. J. (2012) Synthesis and Biological Evaluation of a Biotinylated Paclitaxel With an Extra-Long Chain Spacer Arm. ACS Med Chem Lett 3, 745–748

27. Turkez, H., Tatar, A., Hacimuftuoglu, A., and Ozdemir, E. (2010) Boric acid as a protector against paclitaxel genotoxicity. Acta Biochim Pol 57, 95–97

28. Chatzidarellis, E., Makrilia, N., Giza, L., Georgiadis, E., Alamara, C., and Syrigos, K. N. (2010) Effects of taxane-based chemotherapy on inhibin B and gonadotropins as biomarkers of spermatogenesis. Fertil Steril 94, 558–563

29. Sanchez, A. M., Giorgione, V., Vigano, P., Papaleo, E., Candiani, M., Mangili, G., and Panina-Bordignon, P. (2013) Treatment with anticancer agents induces dysregulation of specific Wnt signaling pathways in human ovarian luteinized granulosa cells in vitro. Toxicol Sci 136, 183–192

30. Ozcelik, B., Turkyilmaz, C., Ozgun, M. T., Serin, I. S., Batukan, C., Ozdamar, S., and Ozturk, A. (2010) Prevention of paclitaxel and cisplatin induced ovarian damage in rats by a gonadotropin-releasing hormone agonist. Fertil Steril 93, 1609–1614

31. Kadota, T., Chikazawa, H., Kondoh, H., Ishikawa, K., Kawano, S., Kuroyanagi, K., Hattori, N., Sakakura, K., Koizumi, S., Hiraiwa, E., and et al. (1994) Toxicity studies of paclitaxel. (I)--Single dose intravenous toxicity in rats. J Toxicol Sci 19 Suppl 1, 1–9

32. Darshan, M. S., Loftus, M. S., Thadani-Mulero, M., Levy, B. P., Escuin, D., Zhou, X. K., Gjyrezi, A., Chanel-Vos, C., Shen, R., Tagawa, S. T., Bander, N. H., Nanus, D. M., and Giannakakou, P. (2011) Taxane-induced blockade to nuclear accumulation of the androgen receptor predicts clinical responses in metastatic prostate cancer. Cancer Res 71, 6019–6029

33. Schiff, P. B., and Horwitz, S. B. (1980) Taxol stabilizes microtubules in mouse fibroblast cells. Proc Natl Acad Sci U S A 77, 1561–1565

34. Oberlies, N. H., and Kroll, D. J. (2004) Camptothecin and taxol: historic achievements in natural products research. J Nat Prod 67, 129–135

