## Supplementary Method for "Oncoprotein GT198 is a direct target of taxol"

### Experimental conditions for GT198 interaction

GT198 is a small homodimeric protein containing a DNA-binding domain. Its DNA-binding function is involved in both DNA repair and transcription. GT198 dimerization is known to be essential for its interaction with nuclear receptors (16). A GT198 dimer may be more physiological in protein conformation. Thus, recombinant His-tagged GT198 was chosen in this study to avoid a large GST tag that might hinder GT198 dimerization in the DNA-binding assays. However, GST-GT198 is capable of binding to biotin-labeled taxol (**Supplementary Figure S2a-b**).

In binding and competition assays, since His-tagged or GST-tagged GT198 proteins are sufficiently pure (**Figure 1d** and **Supplementary Fig. S2a**), the *in vitro* binding between GT198 and taxol represents a direct interaction. In the course of this study, we have found that zinc affects the binding properties of GT198, and that taxol binding to GT198 is zinc-dependent (**Supplementary Figure S2c**). In contrast, zinc significantly decreases the DNA-binding capacity of GT198 (**Supplementary Figure S2d**). Because many DNA repair proteins possess zinc fingers, and GT198 contains histidine and cysteine residues in the DBD (16), GT198 may also contain zinc fingers remained to be further confirmed by structural analysis. Since zinc is present at physiological conditions *in vivo* and in cultured cells, it is likely that the endogenous GT198 protein could adapt a conformation more suitable for taxol interaction than that of the recombinant protein without zinc supplement. Using endogenous GT198 immunoprecipitated from cells, we were able to detect taxol binding without adding zinc (**Supplementary Figure S1b**). Alternatively, zinc could stabilize GT198 structure and render it less accessible for DNA binding. The C terminal domain (aa 180-217) could be potentially involved in such regulation since its deletion improves DNA binding (15). Not shown in this study, we have found GT198 1-180 has increased binding activity than the wildtype GT198 towards other drugs including doxorubicin, which has a competitive interaction with GT198 (23). In this study, green-labeled taxol interaction with GT198 was tested under 1 mM $ZnSO_4$, DNA binding was tested in the absence of zinc, and DNA competition by taxol was tested under 0.1 mM $ZnSO_4$ to maximize taxol interaction while retaining sufficient DNA binding. These observed *in vitro* zinc effects on GT198 potentially reflect the presence of a conformational switch *in vivo* (**Figure 2e**).
